# Time-resolved fluorescence analysis of LHCII in the presence of PsbS at neutral and low pH

**DOI:** 10.1101/456046

**Authors:** Emanuela Crisafi, Maithili Krishnan, Anjali Pandit

## Abstract

In plant chloroplast membranes, non-photochemical quenching (NPQ) is activated as a response to a low luminal pH and controlled by the pH-sensing protein PsbS. It has been proposed that PsbS directly interacts with the light-harvesting complexes (LHCII) of Photosystem II, inducing quenching of LHCII Chl excitations, whilst others proposed that PsbS has an indirect role in controlling the organization of the membrane. In this study, we systematically test the influence of low pH and PsbS on the fluorescence lifetimes of membrane-embedded *spinach* LHCII. The proteoliposome preparations contain LHCII in mild quenched states, aimed to mimic fluorescence conditions of dark-adapted leaves. We find that under those conditions, acidification and the presence of PsbS do not have significant effect on the LHCII Chl fluorescence lifetimes. This supports a view in which the functional role of PsbS consists of re-organizing the thylakoid membrane under light stress, rather than creating direct quencher states. The dimeric form of PsbS appears to be destabilized in lipid membranes compared to detergent micelles, which might explain why the low-pH PsbS crystal structure is dimeric, while *in vivo* activation of PsbS has been correlated with its monomerization at low pH.

**Highlights:**

- PsbS does not affect LHCII Chl excited-state lifetimes
- PsbS dimers have lower stability in lipid membranes than in detergent micelles
- LHCII proteoliposomes with protein-to-lipid molar ratios in the range 0.5.10^−3^ to 1.10^−3^ form suitable membrane models for mimicking dark-adapted fluorescence states

## INTRODUCTION

In natural fluctuating light conditions, plants need to constantly adjust their photosynthetic antenna to the incoming light intensities to prevent themselves from over-excitation and photodamage. They have developed sophisticated feedback mechanisms that regulate light harvesting. In excess light conditions, non-photochemical quenching (NPQ) processes are activated that dissipate a large part of the incoming excitations as heat ^1^.

Light-harvesting complexes (LHCs) that are associated with Photosystem II (PSII) have the ability to reversible switch their conformation into a quenched state, which is assumed to be the cause of excitation quenching during NPQ ^2^. Single-molecule experiments have shown that individual LHCs can fluctuate between fluorescent and quenched states ^3,4^. In bulk experiments, the quenched state is produced upon aggregation of LHCs ^5,6^. Aggregation-dependent quenching has also been observed in native thylakoid membranes, where aggregates of the peripheral light-harvesting complex, LHCII, are formed under highlight conditions ^7,8^.

The fast energy component of NPQ, called qE, in plants is reversibly controlled via the pH-sensing protein Photosystem II subunit S (PsbS), of which the activity is triggered by acidification of the inner compartments of chloroplasts, the thylakoid lumen ^9,10^. Switching back from high to moderate light conditions, PsbS also accelerates fast de-activation of the quenching process ^11^. The fast qE process is connected to a slower quenching process, termed qZ ^12^. In qZ, lumen acidification activates the enzyme violaxanthin de-epoxidase (VDE) to catalyze the conversion of the carotenoid violaxanthin (Vio) to Zeaxanthin (Zea). qZ quenching likely occurs due to the binding of Zea to specific LHCs ^13^. The presence of Zea has been proposed to catalyze qE quenching ^14^ and for Zea-accumulating membranes faster fluorescence induction is observed ^15^.

The membrane protein PsbS is the only member of the LHC multi-gene family that does not bind pigments in specific locations. Instead of acting as a light harvester, this protein solely functions as a pH sensor. PsbS is activated under high-light conditions and its activation is proposed to involve protonation of specific glutamic-acid residues at low lumen pH and monomerization of PsbS dimers ^16^. Several studies have investigated PsbS *in-vivo* interaction partners and showed that PsbS interacts with the LHCII polypeptides (Lhcb1, Lhcb2 and Lhcb3) or with the minor antenna proteins (Lhcb4, Lhcb5 and Lhcb6) under influence of the xanthophyll cycle ^16-20^.

The molecular mechanism for qE excitation quenching has not been resolved and the mechanistic trigger that activates the process via PsbS has not been clarified. Molecular interactions of PsbS with LHCs might induce conformational changes of the antenna proteins into quenched states. PsbS has also been suggested to induce supramolecular rearrangements of LHCs and PSII reaction center complexes ^21^. According to current models, the activation of PsbS at low pH concerns the protonation of two key residues and a monomerization step ^16,22^. Monomerized PsbS may disconnect LHCII from PSII, after which the released LHCII complexes could form clustered aggregates that produce dissipative states via an aggregation-dependent quenching mechanism ^20,23^.

Liposomes form suitable model systems to investigate protein and lipid interactions in membranes of reduced complexity compared to natural biological membranes and dissect the functions of selective membrane components. Various studies have investigated the properties of LHCII proteoliposomes ^4,5,24^ and two studies report on the interaction with PsbS. Liu et al. demonstrated that PsbS could be refolded directly in LHCII-reconstituted membranes ^25^. Wilk et al. co-reconstituted PsbS in liposomes with very low amounts of LHCII to prevent quenching induced by LHCII self-aggregation, and showed that under those conditions the LHCII fluorescence was considerably quenched in the presence of both PsbS and Zea ^26^. The work of Wilk et al. does not report on the effects of acidification, which is known to activate PsbS *in vivo*.

In this study, we perform a systematic fluorescence lifetime analysis on PsbS-LHCII and LHCII-only proteoliposomes at neutral and low pH to gain insight in the molecular origin of the fast qE response that requires PsbS and is activated by low pH before xanthophyll conversion has taken place. For our membrane model, we choose the conditions so that the fluorescence of LHCII-only proteoliposomes at neutral pH mimics the fluorescence of leaves in dark-adapted state. In dark-adapted leaves, the fluorescence of the PSII antenna is moderately quenched and has an average lifetime of ~2 ns, which is considerably shorter than the lifetime of ~4 ns that is observed for isolated LHCII in detergent solutions ^27^, and which suggests that the full fluorescent state of LHCII is not occurring *in vivo*.

## MATERIALS AND METHODS

### LHCII extraction

Light harvesting complexes were purified from spinach leaves (*Spinacia aloracea*) as described in our previous work ^5^. Briefly, LHCII trimer complexes were isolated via sucrose gradient. The trimeric green band of LHCII was extracted using a needle. Purified LHCII complexes were characterized by absorption spectroscopy. The buffer was exchanged into 50 mM HEPES, 100 mM NaCl, pH = 7.5, 0.03% n-Dodecyl *β*-D-maltoside (*β*-DM, purchased from Sigma) and concentrated using Amicon Ultra 2 ml centrifugal filters with cut off of 30 K (Millipore) before to store the protein at –80 °C until use.

### PsbS refolding

Psbs genes from *Physcomitrella patens* and *Spinacia aloracea* were overexpressed in *E. coli* and purified using the protocol from Krishnan et al ^28^. The unfolded PsbS was stored at 4°C until further use. PsbS was refolded in presence of the detergent *β*-DM. According to the refolding protocol, PsbS was mixed with heating buffer (20mM NaOAc, 100mM NaCl, 4% lithium dodecyl sulfate (LDS), 25% sucrose, pH 7.5) and heated to 98°C for 1 minute. Appropriate amount of detergent (β-DM) was added to the mixture. The addition of 200 mM KCl and 30 minutes incubation at 4°C precipitates LDS, thus allowing β-DM to refold PsbS. Aggregated LDS was pelleted by spinning at 20000 x g for 10 minutes at 4°C. The concentration of PsbS protein in mg/ml was determined from SDS page analysis.

### Preparation of asolectin liposomes

Liposomes were prepared according to Crisafi & Pandit with the following adjustments ^5^. Asolectin lipids from soybean (purchased from Sigma) were dissolved in chloroform to a concentration of 5 mg/ml. With a stream on N_2_ the organic solvent was evaporated from the chloroform/asolectin solution collected in a round-bottom flask. To remove all traces of solvent an extra step of evaporation in a rotary evaporator (R3000, Buchi) was performed. The phospholipid bilayer was then hydrated with the desired buffer (50 mM HEPES, 100 mM NaCl, pH 7.5) and the round bottom flask was rotated at low rpm using the rotary evaporator for 1-2 hours. After that if the bilayer was not completely solubilized, the solution was sonicated for 1 min (2210 Branson). The liposome suspension was exposed to 10 freeze/thaw cycles followed by extrusion through polycarbonate membranes of 400 and 200 nm pore size, using a mini extruder (Avanti polar lipids). Sizes of liposome preparations were determined by dynamic light scattering (DLS) using a Malvern Zetasizer Nano ZS equipped with a Peltier controlled thermostatic cell holder.

### Preparation of LHCII and LHCII-PsbS proteoliposomes

For protein insertion, preformed liposomes were destabilized by addition of 0.03% of *β*-DM to facilitate insertion of LHCII and PsbS into the liposome membranes. LHCII complexes were added to the liposome suspension and incubated for 1 hour. Bio beads were added to the suspensions in 2 steps. The first incubation step was performed at 4°C overnight while gently rotating the sample using a roller mixer SRT9D (Stuart). Subsequently, the Bio beads were refreshed and the solution was again incubated for 2-3 hours. The proteoliposome suspensions were centrifuged at 20000g for 10 min at 4 °C using a table-top centrifuge (3K18, Sigma) to remove non-incorporated LHCII and PsbS that forms aggregate pellets 29. LHCII insertion and removal of LHCII aggregate pellet was confirmed by running LHCII and LHCII-PsbS proteoliposomes over a sucrose gradient that showed a single green band containing the proteoliposomes ^30^. The reported protein to lipid ratios (PLR) are defined as moles of LHCII or PsbS trimers per moles of total lipids. The concentration of LHCII trimers was determined from the molar extinction coefficient for LHCII (trimers) at 670 nm, ε = 1,638,000 M^−1^ cm^−1 31^.

### Circular Dichroism (CD) experiments

CD spectra were recorded on a J-815 spectrometer (Jasco, Gross- Umstadt,) equipped with a temperature control set. The wavelength range was from 350 to 750 nm, data pitch 1 nm, response 2 s, band width 2 nm and scanning speed of 50 nm/min at 20 °C using a 1 cm quartz cuvette (Hellma).

### UV/VIS absorbance experiments

Absorption spectra were recorded on a UV-1700 PharmaSpec UV–visible spectrophotometer (Shimadzu,) with the wavelength range set from 350 to 750 nm using 1×1cm cuvette.

### Steady-state fluorescence experiments

Fluorescence measurements were performed on a Cary Eclipse fluorescence spectrophotometer (Varian), collecting emission spectra from 660 to 720 nm using 3×3 mm quartz cuvettes. The optical density of the sample preparations varied from 0.03 to 0.07 cm^−1^ at 650 nm. The excitation wavelength was set at 650 nm or 475 nm.

### Time resolved fluorescence experiments

Time-resolved fluorescence measurements were performed using a FluoTime 300 (PicoQuant) time-correlated photon counter spectrometer. Samples were hold in a 1×1 cm quartz cuvette that was thermostated at 20 °C and excited at 440 nm using a diode laser (PicoQuant). Fluorescence decay traces were fitted with multi-exponentials using a χ^2^ least-square fitting procedure.

### Gel electrophoresis

For Sodium Dodecyl Sulphate PolyAcrylamide Gel Electrophoresis (SDS-page gel) analysis, proteoliposomes were pelleted and re-suspended in HEPES buffer and were analyzed on 12,5% polyacrylamide running gel with 4% stacking gel. Gels were stained using Silver Stain Plus kit (Bio-Rad) according to the standard Bio Rad protocol. Molecular masses were estimated via a molecular standard (precision plus protein dual color, Bio-Rad).

## RESULTS

### Relationship between the LHCII protein-to-lipid ratio (PLR) and fluorescence quenching

LHCII is known to form fluorescence-quenched states in its aggregated form. To separate fluorescence effects induced by PsbS and acidification from the effects of LHCII self-aggregation, we first aim to have an overview of how quenching of LHCII in liposomes depends on the molar protein to lipid ratios (PLR), focusing on the regime where LHCII proteoliposomes display moderate fluorescence. Our previous work reported on the fluorescence yields of LHCII proteoliposomes at very low LHCII concentrations, and a single-molecule fluorescence study reported on the fluorescence properties of individual proteoliposomes in this range ^4,5^. In addition, the work of Wilk et al. used LHCII-proteoliposomes with dilute LHCII concentrations ^26^. We decided to combine data results from the different studies in one graph (Fig. 1), where quenching of LHCII proteoliposomes is plotted against the PLR. To generate this plot, the fluorescence yields were calculated by dividing the reported average fluorescence lifetimes or yields of LHCII proteoliposomes by the values for LHCII in detergent solution reported in the same studies. As shown in Fig. 1, data from different sources follow a remarkably similar trend, considering that different preparation techniques were used and that our previous study used asolectin proteoliposomes, whilst the other studies used liposomes prepared from thylakoid lipid mixtures. At very low protein densities, a steep depcrease of the fluorescence is observed with increased LHCII concentrations. In this PLR regime, the liposomes contain only a few proteins per vesicle according to single-molecule spectroscopy and electron freeze-fracture microscopy ^4,26^. At higher PLRs there is a more gradual increase of quenching with increased LHCII concentrations. The data in Fig. 1 could not be fitted properly with a single exponential curve and is fitted with two exponentials. Fluorescence data of LHCII in lipid nanodiscs are also plotted in the graph. LHCII complexes that are captured in nanodiscs are prevented from self-aggregation. Under those conditions no significant quenching occurs, demonstrating that quenching is not induced by protein-lipid interactions *per se* ^4,5^. Because the work on highly diluted LHCII proteoliposomes report that clusters of only few LHCIIs are required to trigger quenching, we assume that the steep slope in the curve in Fig. 1 at low PLRs reflects the transition going from single to multiple LHCIIs per vesicle.

**FIGURE 1.**
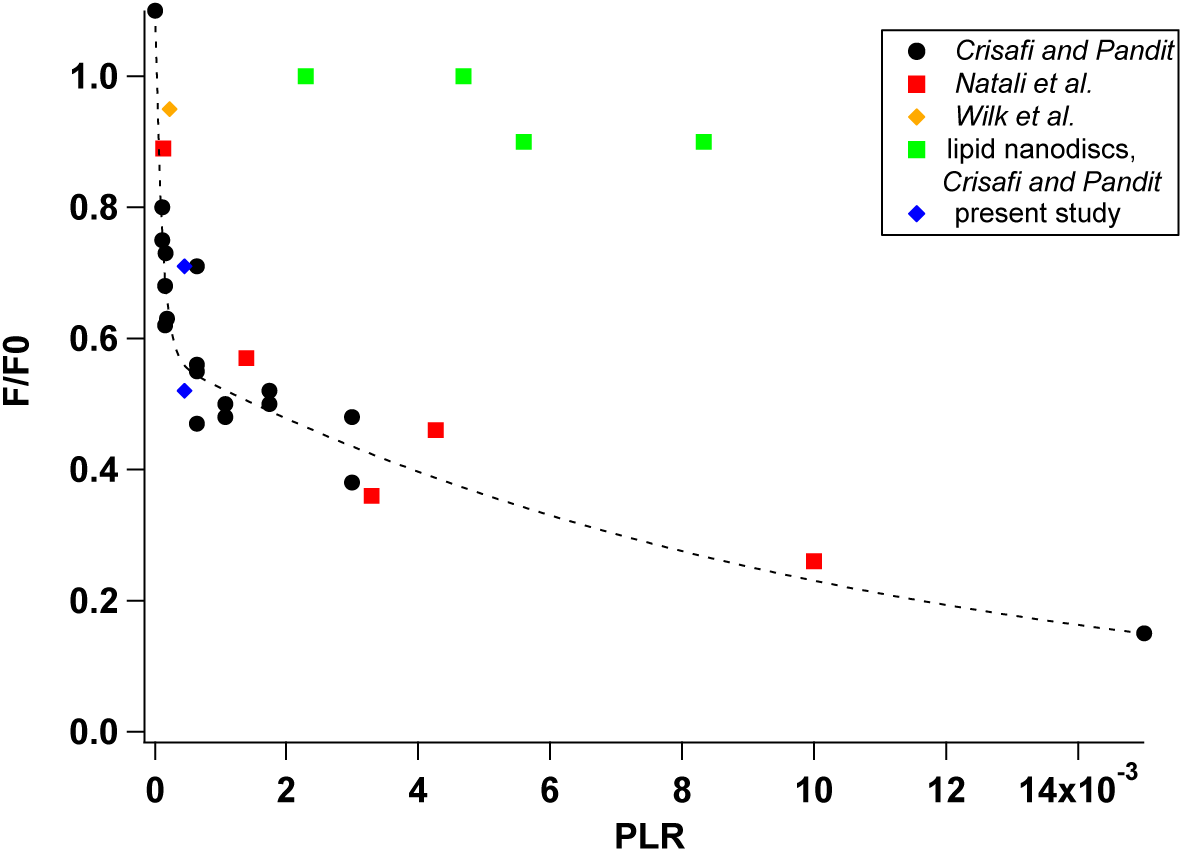
Fluorescence yield of LHCII proteoliposomes and nanodiscs versus the protein to lipid ratio. Data results from three different studies is plotted. *Crisafi and Pandit* ^5^ used liposomes prepared from asolectin, while *Natali et al.* ^4^ and *Wilk et al.* ^26^ used mixtures of galactolipids. Proteoliposome data results are fit with a double exponential fit.

The fluorescence yield of 50% around the breakpoint in the curve in Fig. 1 corresponds to an average fluorescence lifetime of ~2 ns, which resembles the fluorescence lifetimes of dark-adapted leaves with closed reaction centers ^15^. With higher protein concentrations, the fluorescence yield gradually decreases, approaching the fluorescence characteristics of leaves in light-adapted, photoprotective states for which lifetimes of 0.3 to 0.5 ns have been reported ^15^.

For our proteoliposome preparations, we choose to use a PLR of 6.5*10^−4^ (LHCII:lipid, *mol*/*mol*), which is close to the breakpoint in the slope of the curve, in order to mimic the natural fluorescence conditions of antennas in dark-adapted leaves. Protein densities in native thylakoid membranes are much higher than in our proteoliposomes and we do not intend to also reproduce *in vivo* protein densities, as this would lead to formation of large LHCII aggregates that are strongly quenched. In native membranes, involvement of LHCII in super-complexes may prevent formation of large LHCII aggregates.

### Analysis of PsbS proteoliposomes

Proteoliposomes are prepared from asolectin lipids, of which we have better control over the protein insertion than for lipid mixtures that mimic the natural lipid compositions of plant thylakoid membranes. The natural lipid composition of plant thylakoid membranes involves high amounts (~40%) of the nonbilayer lipid monogalactosyl diacylglycerol (MGDG) that tends to form inverted hexagonal phases. MGDG lipids can be incorporated in lipid bilayers of proteoliposomes with high protein densities, but preparing diluted proteoliposomes with high amounts of MGDG is challenging and leads to unstable vesicles. As a result, protein insertion yields tend to vary, which complicates quantitative comparison between different samples. As shown in Fig. 1, the trend for aggregation-dependent quenching of LHCII in asolectin liposomes ^5^ is similar to the trend for liposomes prepared from a mixture of thylakoid lipids ^5^. For co-reconstitution, a PsbS:LHCII molar ratio of 1:1 is used. This ratio is higher than the natural occurrence of PsbS in thylakoid membranes and is used to increase the probability of PsbS-LHCII interactions.

Insertion of LHCII in the liposomes is confirmed by the absence of LHCII aggregate pellet in reconstitution solutions and by a sucrose gradient analysis that shows a green band containing the LHCII proteoliposomes. To test the insertion of PsbS, which is not visible in the sucrose gradient, we perform an SDS-page gel analysis on PsbS-only proteoliposomes. The use of Western blotting was discarded as we notice that PsbS dimers can give a false-positive reaction in anti-Lhcb1 Western blots (see Fig. S1).

PsbS forms a monomer-dimer equilibrium in detergent solutions ^28^. Aging of the protein solutions shifts the equilibrium towards the dimeric form, which proceeds more slowly at low pH. To check if the initial state of PsbS influences the dimerization state inside membranes, PsbS insertion in preformed liposomes is tested under four different conditions. Proteoliposomes are prepared at pH 5.0 or pH 7.5 and two PsbS preparations are used for insertion that contain PsbS in different monomer-dimer equilibria. The first PsbS sample is three days old and contains more dimers than monomers according to SDS-page analysis (Fig. 2a). The second PsbS sample is freshly prepared and contains more monomers than dimers. The results of our insertion tests confirm that PsbS inserts into the liposomes under all four conditions (Fig. 2b). The PsbS fractions run only as a monomer band in the SDS-page gel, irrespective of the pH conditions or oligomeric state before insertion. When the insertion is performed in pH5.0 buffers, however, protein pellets is observed in the liposome suspensions and the insertion appears to be less efficient. For this reason we proceed with the reconstitution of PsbS in pH7.5 buffer solutions. From a comparison of the gel band intensities of PsbS proteoliposomes to those of the removed pellet, we estimate that at pH 7.5 ~80% of the PsbS is inserted in the liposomes (Fig. S2). SDS-page gel analysis of PsbS-LHCII proteoliposomes confirms that both proteins are co-inserted (Fig. S3).

**FIGURE 2.**
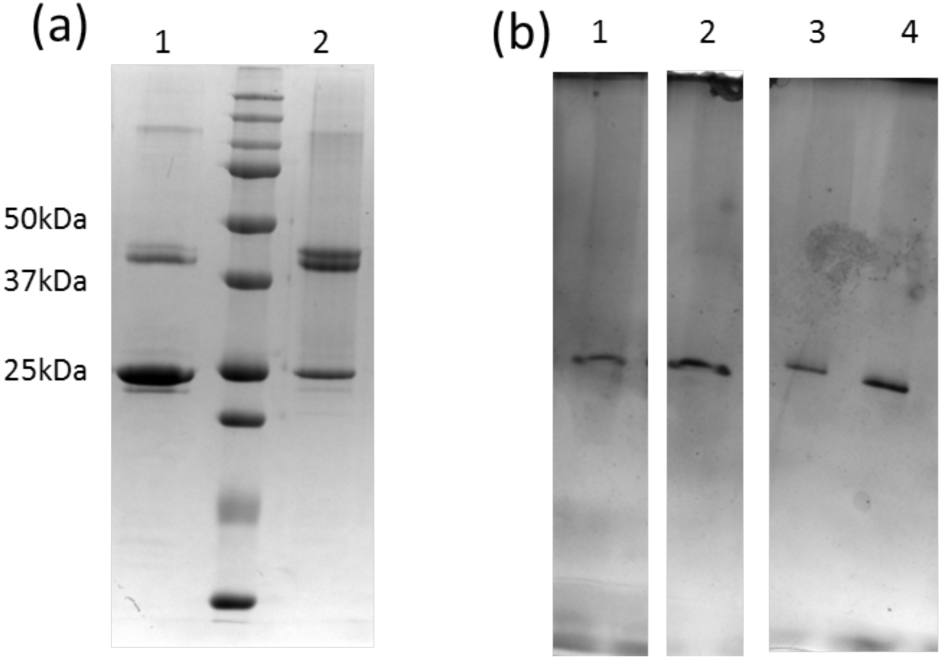
(a) SDS-page gel of PsbS in detergent solution before liposome insertion. Sample 1 contains PsbS that was predominantly in the monomeric form and 2 contains PsbS that is predominantly in the dimeric form. The middle lane contains the protein marker. (b) PsbS proteoliposomes confirming PsbS insertion (silver stain gel). Lane 1 and 2: proteoliposomes prepared at pH 5.0 using sample 1 (1) or sample 2 (2). Lane 3 and 4: proteoliposomes prepared at pH 7.5 using sample 1 (3) or sample 2 (4).

### Fluorescence lifetimes analysis of LHCII-PsbS proteoliposomes at neutral and at low pH

Time-resolved fluorescence experiments are performed on LHCII-only and LHCII-PsbS proteoliposomes that are equilibrated at pH 7.5 or at pH 5.0. Presented LHCII-only and LHCII-PsbS proteoliposome fluorescence data that are compared were always prepared from the same liposome batches, since some variation in the fluorescence lifetimes is found from batch to batch. To equilibrate the proteoliposome preparations at pH 5.0, we use a protocol in which suspensions prepared at pH 7.5 are acidified by injection of HCl followed by the addition of nigericin, a liposoluble compound, in order to equilibrate pH conditions inside and outside the proteoliposomes ^32^. Dynamic Light Scattering (DLS) measurements on liposomes before and after acidification and nigericin treatments confirm that there are no significant changes in the diameter sizes of the proteoliposomes (Table S1). Upon acidification, the contribution of very large particles (>500 nm) disappears in the DLS size distribution diagrams and therefore the average diameter is slightly shifted to lower values.

Table 1 presents the results of the fluorescence lifetime experiments that are performed on LHCII-only and LHCII -*patens* PsbS proteoliposomes. For comparison, the fluorescence lifetimes of LHCII in 0.03% *β*-DM detergent solutions equilibrated at pH 7.5 or at pH 5.0 are also shown. The presented results are the average of two reconstitution experiments using the same batch of preformed liposomes. The results show that acidification causes at most ~4% reduction of the average fluorescence lifetimes, whilst no significant differences are observed comparing LHCII and LHCII-PsbS proteoliposomes. Steady-state fluorescence spectra collected with 440 nm excitation show spectral broadening of the emission peak and a red shoulder around 700 nm for the proteoliposomes compared to LHCII in *β*-DM (Fig. 3). The two characteristic effects are more pronounced for PsbS-containing proteoliposomes while no changes are observed at low pH.

**Table 1.** Fluorescence lifetimes of proteoliposomes using a PLR of 4.65·10^−4^. PsbS containing proteoliposomes contained 1:1 (*mo*l PsbS/*mol* LHCII trimer)p*Patens* PsbS.

| Sample | A1% | $\tau_1$ | A2% | $\tau_2$ | A3% | $\tau_3$ | $\tau_{av}$ |
| --- | --- | --- | --- | --- | --- | --- | --- |
| **LHCII proteolip pH 7.5** | 28.4 | 4.4 | 52.7 | 2.6 | 18.9 | 0.8 | 2.7 |
| **LHCII proteolip pH 5.0** | 21.9 | 4.5 | 56.0 | 2.5 | 22.1 | 0.8 | 2.6 |
| **LHCII+PsbS proteolip pH 7.5** | 21.5 | 4.6 | 58.0 | 2.6 | 20.4 | 1.0 | 2.7 |
| **LHCII+PsbS proteolip pH 5.0** | 19.4 | 4.6 | 57.8 | 2.5 | 22.8 | 1.0 | 2.6 |
| **LHCII in β-DM pH 7.5** | 72.1 | 4.2 | 27.9 | 2.9 | - | - | 3.8 |
| **LHCII in β-DM pH 5.0** | 61.4 | 4.3 | 35.3 | 2.8 | 3.3 | 1.1 | 3.7 |

**FIGURE 3.**
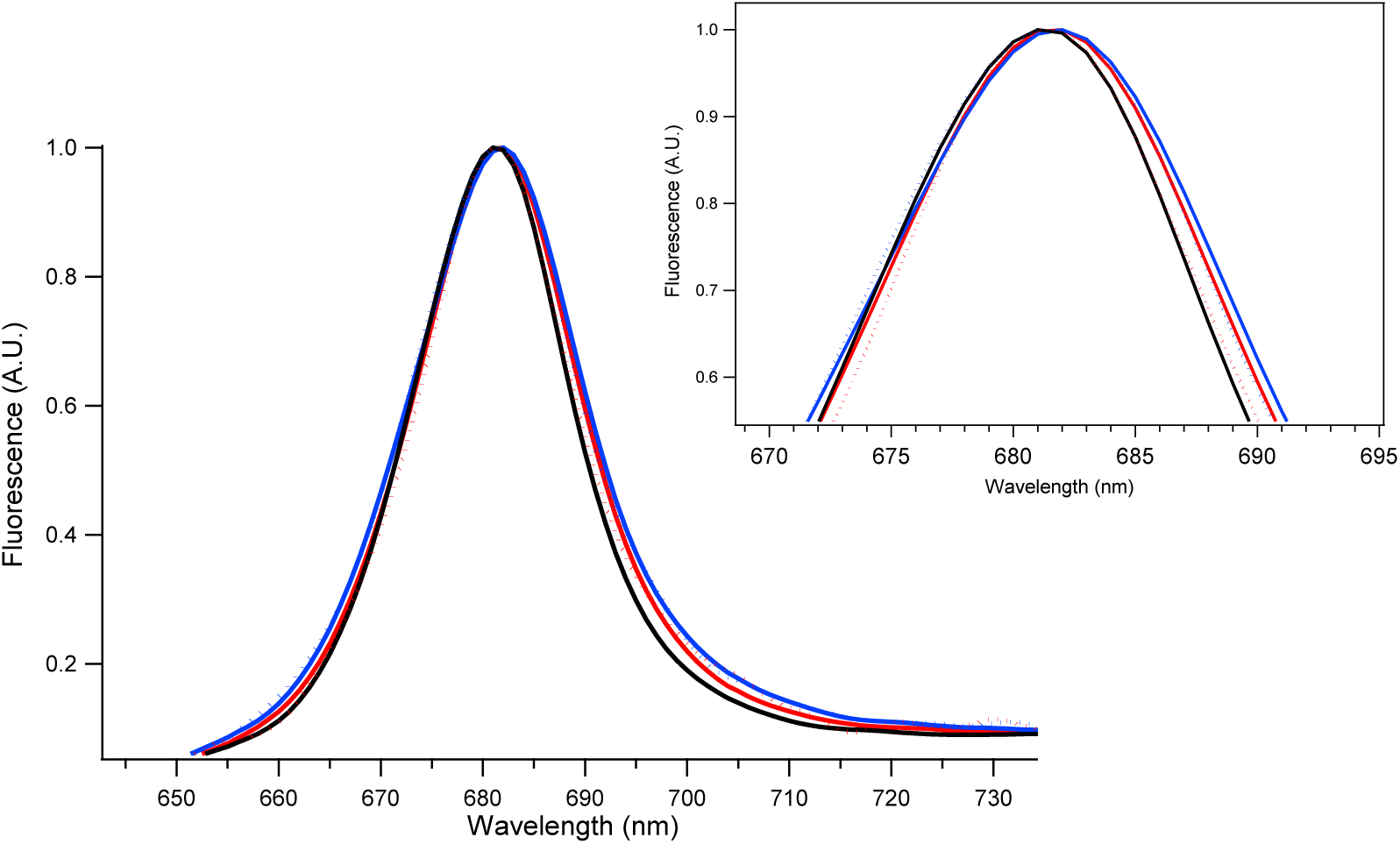
Fluorescence emission spectra of LHCII in *b*-DM (black), LHCII-only proteoliposomes (red) and LHCII-PsbS proteoliposomes (blue) at pH7.5 normalized at the fluorescence maximum. Dashed lines: LHCII only proteoliposomes (red) and LHCII-PsbS proteoliposomes (blue) at pH5.0.

In the above experiments, we used PsbS from the moss *Physcomitrella patens* because an extensive analysis of this protein has been performed ^28^, and we reconstituted *patens* PsbS together with LHCII extracted from spinach leaves. Since no significant effect of PsbS is observed, we consider that functional interactions between PsbS and LHCII may rely on molecular recognition sites that require protein interaction partners from the same organism. Therefore experiments are repeated on PsbS-LHCII proteoliposomes that contain PsbS from *Spinacia oleracea.* Fig. 4 and 5 compares the results of a fluorescence analysis on PsbS-LHCII liposomes containing *patens* PsbS (a) or *spinach* PsbS (b). The data results show that the lifetime distributions of proteoliposomes containing moss or spinach PsbS are very similar and that in both cases, the fluorescence lifetimes are not significantly affected by acidification.

**FIGURE 4.**
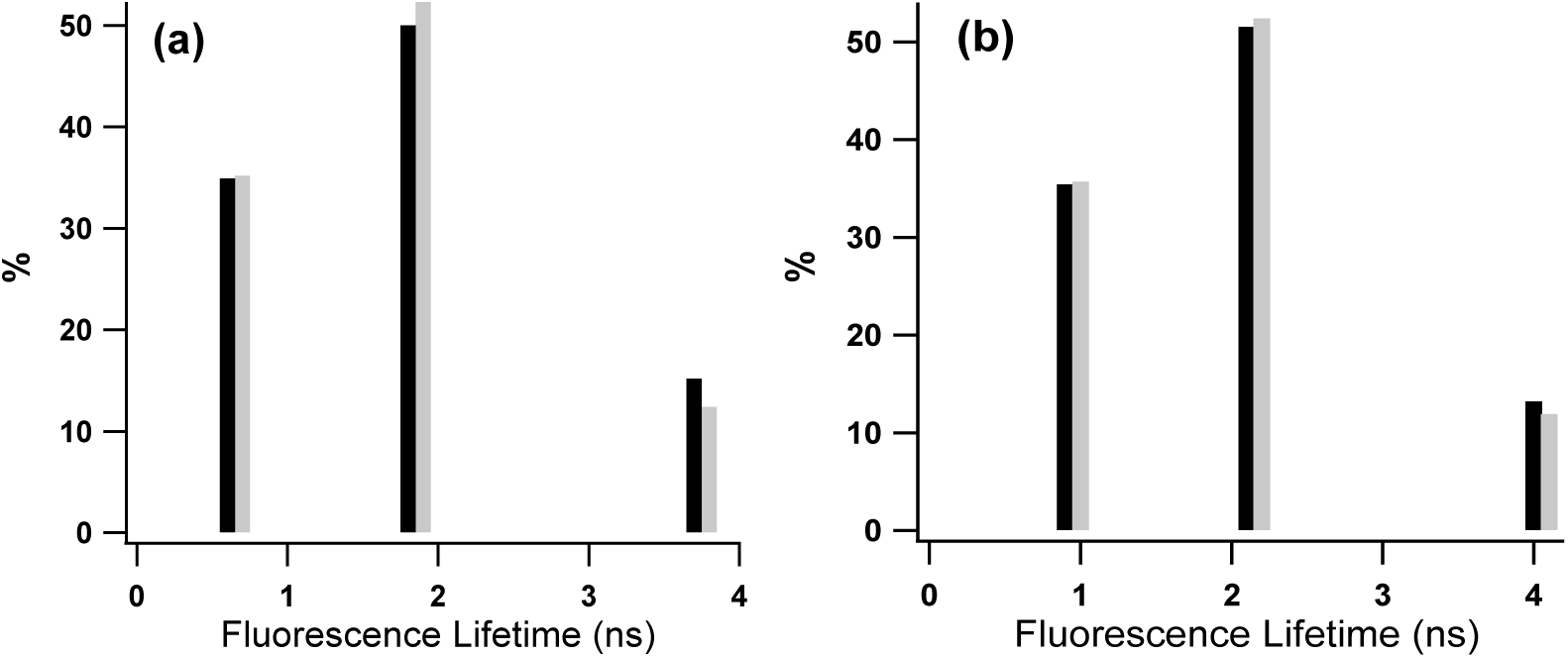
Time-resolved fluorescence analysis of PsbS-LHCII proteoliposomes containing *patens* PsbS (a) or *spinach* PsbS (b). Black sticks: pH 7.5 conditions, grey sticks: pH 5.0 conditions.

**FIGURE 5.**
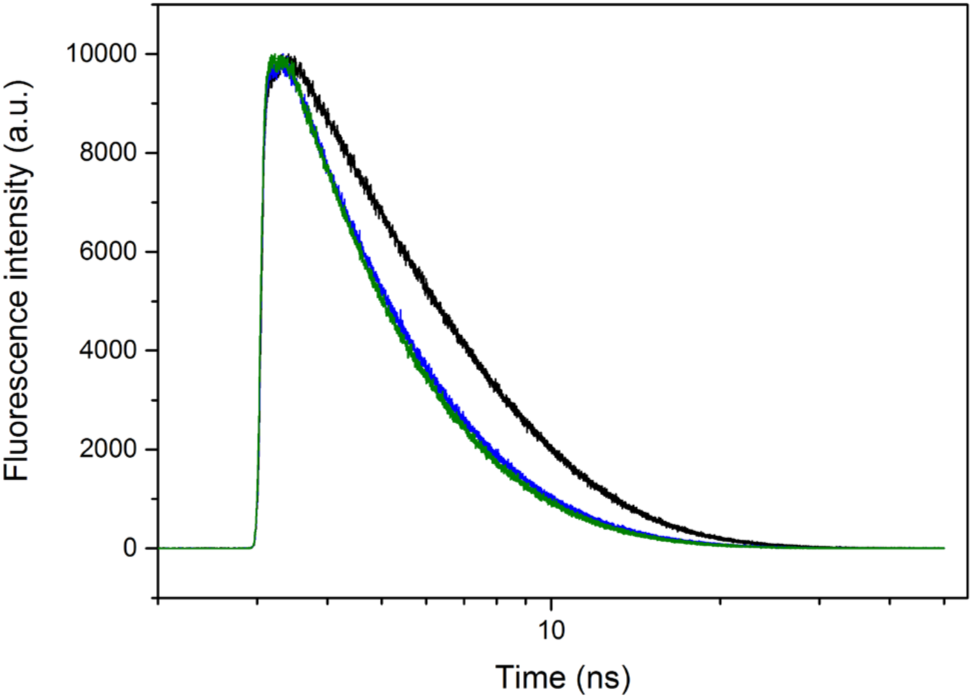
TRF experimental and fit traces of the data presented in Fig.3 (b). Black: LHCII in 0.03% β–DM, blue: LHCII + PsbS proteoliposomes at pH7.5, green: LHCII+ PsbS proteoliposomes at pH 5.0.

### CD spectral analysis of LHCII-PsbS proteoliposomes

The pigment excitonic CD spectrum of LHCII is very sensitive to the changes in the protein conformation or in the protein micro-environment. The excitonic CD spectra in the visible region are collected of LHCII-only and of LHCII-PsbS proteoliposomes and presented in Fig. 6. PsbS does not bind any pigments and the CD bands originate from the excitonic interactions among the LHCII Chl and carotenoid pigments. The low-pH CD spectra of PsbS-only and PsbS-LHCII proteoliposomes are very similar exept for the red bands at 650 and 680 nm that have somewhat increased intensity for the PsbS-LHCII proteoliposomes while the band at 665 nm has reduced intensity. At low pH, an increase of the negative bands at 450 nm and 461 nm is observed. The CD spectrum of LHCII in β-DM detergent is also plotted. The difference comparing the CD spectral shape of LHCII in β-DM and LHCII proteoliposomes is characteristic for the exchange from a detergent to lipid environment ^5^.

**FIGURE 6.**
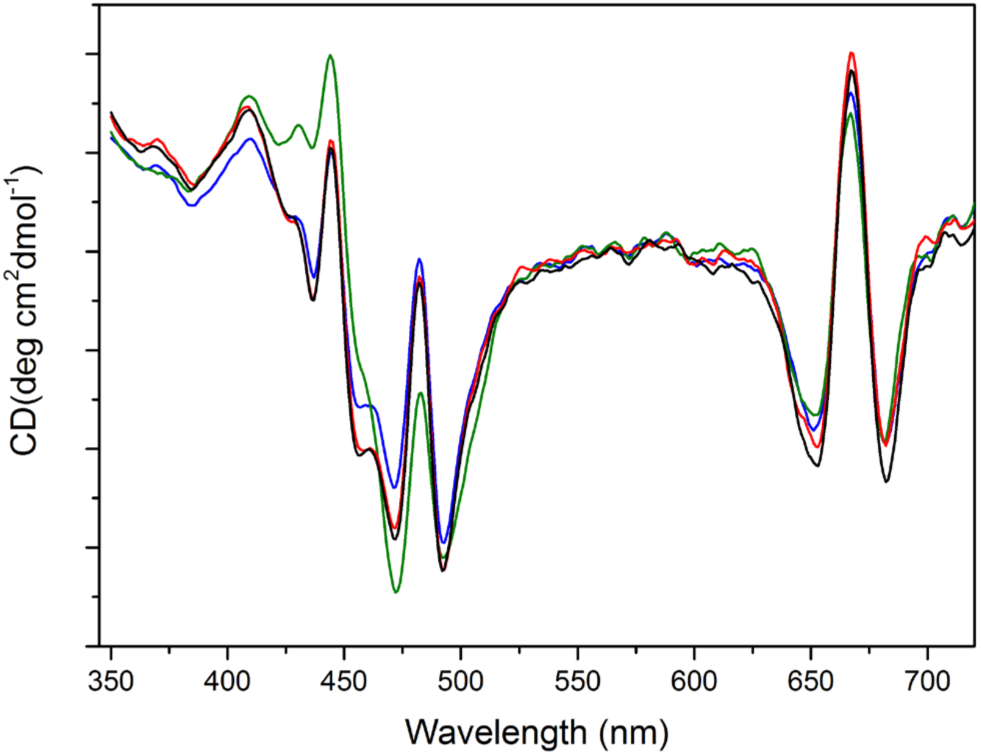
CD spectra of LHCII in 0.03% βDM at pH7.5 (green), LHCII*-*PsbS proteoliposomes at pH 7.5 (blue, fluorescence τ_av=_ 2 ns), LHCII proteoliposomes at pH 5.0 (red, fluorescence τ_av=_ 1.6 ns) and LHCII-PsbS proteoliposomes at pH 5.0 (black, fluorescence τ_av=_ 1.7 ns).

Acidification of LHCII in β-DM detergent does not influence the CD spectral shape (Fig. S5) indicating that the observed difference comparing the CD spectrum at pH 7.5 and at pH 5.0 is not due to conformational changes in individual LHCIIs. The proteoliposome CD samples at neutral and low pH conditions were obtained from different preparations that have small variations in their fluorescence lifetimes. The pH 5.0 CD samples have shorter average lifetimes (1.6 and 1.7 ns for LHCII-only and for LHCII-PsbS proteoliposomes respectively) than the pH 7.5 sample (2.0 ns). We suspect that the small increase in quenching has an effect on the CD spectral shape, which suggests that CD spectroscopy is a very sensitive tool for detecting quenching-related structural changes.

## DISCUSSION

The small increase in the red shoulder and broadening of the fluorescence emission spectrum of PsbS-LHCII versus LHCII-only proteoliposomes indicates that PsbS influences the LHCII fluorescence properties and suggests that PsbS and LHCII proteins are in physical contact with eachother in the lipid membranes. The fluorescence lifetime analysis shows that the presence of PsbS does not have an effect on the LHCII Chl excited-state lifetimes. With our model, we aimed to reproduce the fast NPQ activation mechanism and we did not incorporate Zea that is responsible for the slower contribution of NPQ and depends upon the activation of VDE enzymes. Wilk et al. performed experiments on PsbS-LHCII proteoliposomes in Zea-containing membranes ^26^. On basis of Western blotting experiments it was concluded that PsbS-LHCII heterodimers were formed, and it was shown that LHCIIs in PsbS and Zea-containing proteoliposomes were quenched via carotenoid-Chl interactions. The authors infer that the combination of PsbS and Zea reduces LHCII fluorescence yields via one-to-one interactions between LHCII and PsbS. We found that PsbS reacts to Lhcb1 antibodies and warn that their conclusions from Western blotting could be based on a false-positive result. Nevertheless, their data suggest that the interplay between Zea, PsbS and LHCII could play a role in Zea-regulated quenching. A difference between their sample conditions and ours is that they used LHCII proteoliposomes with very low protein densities that contained LHCII in a fully fluorescent state, while we use LHCII proteoliposomes with higher protein densities in which the LHCIIs already have reduced fluorescence without the presence of PsbS. We are not able to reproduce the result of Liu et al. ^25^, who detected a drop in the fluorescence intensity of PsbS-LHCII proteoliposomes upon acidification. We also monitored the fluorescence intensity during acidification and did not observe notable changes (*data not shown*), consistent with the fact that the fluorescence lifetimes do not change at low pH. The approach of Liu et al. consisted of direct refolding of PsbS in preformed liposome membranes. We tested this method and indeed could achieve successful refolding of PsbS directly into asolectin liposomes. Because inspection by eye showed the presence of foam remaining in the liposome suspensions, we suspect that in our hands not all of the LDS detergent was removed with this procedure and we continued with reconstitution starting from refolded PsbS in detergent micelles as described in the Mat and Methods.

In low-detergent solutions, lowering of the pH significantly enhances fluorescence quenching of assemblies of LHCII and this concept has been used to mimic the photoprotective conditions of LHC proteins *in vitro* ^33,34^. We show here that acidification does not shorten the fluorescence lifetimes of LHCII assemblies in proteoliposomes, indicating that LHCII assemblies in the native environment of a lipid membrane are not responsive to lowering of the pH. As shown in Fig. S5, acidification does not induce the quenched state of isolated LHCIIs and the reported increased fluorescence quenching of LHCs low-detergent solutions upon acidification is likely due to enhanced aggregation. In solutions, membrane proteins are capable of assembly into three-dimensional aggregates, whilst in liposomes only lateral protein interactions occur and aggregate sizes are limited to the number of LHCIIs per vesicle. The limitations imposed by the liposome vesicle dimensions may prevent increase of aggregation and enhanced quenching at low pH.

The protein PsbS forms equilibrium between monomers and dimers in detergent solutions. The equilibrium tends to shift towards dimers over time, which occurs faster at neutral pH (^28^ and Fig. 2). It is therefore surprpising to observe that fresh preparations of PsbS proteoliposomes only contain PsbS monomers according to the SDS-page gel analysis. Because denaturing gel conditions were used, PsbS dimers may have formed inside the membranes that did not resist the denaturation conditions. The results either way implicate that PsbS dimers are less stable in lipid membranes than in detergent micelles. Lipids and detergents are both capable of solubilizing membrane proteins through shielding of the protein hydrophobic sites from the aqueous environment. However, while lipid bilayers have a well-defined hydrophobic dimension, this is not the case for detergent micelles and the difference in environment can affect protein local structure ^35^. For PsbS, the two active glutamates that are responsive to pH are located at the membrane-water interface and their direct environment could influence the stability of PsbS dimers. The importance of the micro-environment for PsbS stability could explain why the low-pH crystal structure of PsbS is a dimer, while its low-pH activation *in vivo* has been correlated with monomerization ^10,22^.

The results show that under conditions where LHCII is in a mild quenched state, *in vitro* co-insertion of PsbS does not affect the excited-state lifetimes of LHCIIs, discarding qE models that are based on pH-induced PsbS-LHCII molecular interactions. The strong dependence of the fluorescence of LHCII proteoliposomes on the LHCII/lipid ratios, combined with challenges of protein-membrane reconstitution and reproducibility that we experienced, especially for PsbS that is prone to strong aggregation, makes that membrane reconstitution studies on this class of proteins should be carefully conducted. In that respect, LHCII proteoliposomes in moderate quenched states form easier controllable models than highly diluted proteoliposomes, in which small changes in the number of LHCIIs per vesicle will significantly influence the fluorescence states.

While we did not observe a significant effect of PsbS on the fluorescence states of LHCII, the strong effect of LHCII-LHCII interactions on their fluorescence properties is well known. Aggregation-dependent fluorescence quenching of LHCII in membranes was first reported by Moya et al. and the relationship between LHCII cluster sizes and quenching has been analyzed in detail using single molecule spectroscopy, lipid nanodiscs, and combined FLIM/AFM on planar lipid membranes ^4,5,24,36^. Fig. 1 shows that starting from conditions that mimic dark-adapted states in proteoliposomes with only few LHCIIs (i.e. proteoliposomes with a fluorescence yield of ~50% compared to LHCII in detergent), qE-mimicking conditions (a fluorescence yield of ~15%) can be achieved by a 7-10 fold increase of the LHCII densities. Thus, in absence of additional quenching mechanisms, a membrane response that creates antenna rearrangements going from a few LHCIIs to 20-30 connected LHCIIs would be sufficient to drive a transition from dark-adapted states towards qE states. Our results suggest that the pH-dependent role of PsbS during the fast qE response lies in creating membrane rearrangements and super-complex remodeling ^30,37^, which may facilitate LHCII aggregation quenching, rather than in creating direct quencher states.

## ACKNOWLEDGEMENTS

This work was financially supported by a CW-VIDI grant of the Netherlands Organization of Scientific Research (NWO) under grant nr. 723.012.103.

